# A New Behavioral Paradigm for Frustrative Non-reward Reveals a Global Change in Brain Networks by Frustration

**DOI:** 10.1101/2023.02.28.530477

**Authors:** Aijaz Ahmad Naik, Xiaoyu Ma, Maxime Munyeshyaka, Ellen Leibenluft, Zheng Li

## Abstract

**Background:** Irritability, defined as proneness to anger, can reach a pathological extent. It is a defining symptom of Disruptive Mood Dysregulation Disorder (DMDD) and one of the most common reasons youth presents for psychiatric evaluation and care. Aberrant responses to frustrative non-reward (FNR, the response to omission of expected reward) are central to the pathophysiology of irritability. FNR is a translational construct to study irritability across species. The development of preclinical FNR models would advance mechanistic studies of the important and relatively understudied clinical phenomenon of irritability.

**Methods:** We used FNR as a conceptual framework to develop a novel mouse behavioral paradigm named Alternate Poking Reward Omission (APRO). After APRO, mice were examined with a battery of behavioral tests and processed for whole brain c-Fos imaging. FNR increases locomotion and aggression in mice regardless of sex. These behavioral changes resemble the symptoms observed in youth with severe irritability. There is no change in anxiety-like, depression-like, or non-aggressive social behaviors. FNR increases c-Fos+ neurons in 13 subregions of thalamus, iso-cortex and hippocampus including the prelimbic, ACC, hippocampus, dorsal thalamus, cuneiform nucleus, pons, and pallidum areas. FNR also shifts the brain network towards a more global processing mode.

**Conclusion:** Our novel FNR paradigm produces a frustration effect and alters brain processing in ways resembling the symptoms and brain network reconfiguration observed in youth with severe irritability. The novel behavioral paradigm and identified brain regions lay the groundwork for further mechanistic studies of frustration and irritability in rodents.

## Introduction

Irritability, defined as proneness to anger, can reach a pathological extent and is one of the most common reasons for psychiatric evaluation and treatment in youth (1). It is a hallmark symptom of Disruptive Mood Dysregulation Disorder (DMDD), a diagnosis introduced into the Diagnostic and Statistical Manual of Mental Disorders (DSM-5) in 2013. Irritability is associated with long-lasting adverse outcomes including high rates of school suspensions, hospitalizations, suicidality, and diagnoses of anxiety and depression (2, 3). Current treatments for irritability are limited and not specific (4). Increased understanding of its neurobiological mechanisms is warranted to facilitate the development of novel, specific interventions. However, despite increased irritability research over the past decade, the neuroscience of irritability is still in its infancy (5–8). Progress has been slow partly due to the lack of behavioral paradigms for studying irritability in model organisms.

Youth with severe irritability have elevated responses to frustrative non-reward (FNR) i.e., the emotional and behavioral response to the omission of an expected reward (9). FNR is a translational, cross-species construct that can be leveraged to uncover pathophysiological mechanisms of irritability. American psychologist Abram Amsel, who pioneered the study of FNR, proposed that instrumental behavior is learned through frustrating, rewarding, and punishing events (10–13). In contrast to the much-studied rewarding and punishing events, little research has focused on the neural substrates of frustrating events. Amsel studied the effect of FNR on behavior using a double-runway setup in which rats were trained to pass two runways connected in series to receive rewards at the end of each runway. After rats learned the task, FNR was introduced by reward omission (10, 13). Amsel demonstrated that FNR invigorates behavior as indicated by faster running and increased resistance to extinction.

Besides the double-runway paradigm, an operant conditioning chamber has also been used to produce frustration in rodents. This method works better for adult animals with well-developed muscles than for juveniles because animals must press a lever repeatedly to receive reward. Both the double-runway and operant conditioning chamber methods, however, require weeks of training (14–16). Long training periods are unsuited to study irritability-like behavior in mice, as DMDD is a disorder diagnosed in children aged 6–18 years, which corresponds to 3–7 weeks of ages in mice (17). The short developmental period of mice restrains the duration of the behavioral paradigm that can be used.

Human studies of brain-based mechanisms of irritability are limited and have yielded mixed results (18). Preliminary studies find that high levels of irritability following frustration in people with DMDD correspond with increased frontal-striatal activation in the anterior cingulate cortex, dorsolateral prefrontal cortex, caudate, amygdala, and parietal cortex (9, 19, 20). Recent network-based pilot studies using both task-based and resting-state fMRI found predictive associations between irritability and post-frustration connectivity in limbic, reward, and sensorimotor networks (9, 21). However, human fMRI studies cannot identify the impact of frustration at the level of neuronal ensembles due to inadequate spatial and temporal resolutions. The capacity to do so in animals would be a significant translational advance that could eventually lead to more precisely tailored innovative treatments for irritability and DMDD.

To study FNR in young mice, we developed a new behavioral paradigm named Alternate Poking Reward Omission (APRO). We show that after exposing juvenile mice to FNR using APRO, mice increase locomotion, aggression, and resistance to extinction of instrumental behavior. Whole brain mapping of neural activation shows that 13 brain regions are activated, and the processing mode of brain network is changed by FNR.

## Materials and Methods

### Alternate poking reward omission (APRO)

Mice (35 days old; male and female) were placed under water restriction for three days when they were given ad libitum access to water for 1 hour per day. The apparatus used for APRO was constructed by NIMH Section on Instrumentation. It is a closed running track (50L x 10W x 15H cm) with a spout installed at each end and a LED light mounted next to the spout. The mouse was allowed to run freely in the track for one 15-min session per day. The mouse could poke the port when it reached the end of the track. If the mouse poked the port other than the previous one it had poked, the LED light came on for 2 sec and a water drop was delivered from the spout.

During training sessions, the mouse received water reward for every correct port poking. Only mice making ≥12 licks on each side by Day 3 proceeded to Day 4 and 5 of APRO. On Days 4 and 5, mice run on the same track with the same rules. While control mice received water reward for all correct pokings on Days 4 and 5, water reward was delivered for only 50% of correct pokings on Day 4 and 20% on Day 5 for mice subjected to FNR. Thirty minutes after the FNR session on Day 5, the mice were subjected to a battery of behavioral tests, including the open field, resident intruder, elevated zero-maze, light/dark box, forced swim, sucrose preference, and three-chamber social preference tests. Different cohorts of mice were used for each behavioral test so that each mouse was subjected to only one behavioral test. For the extinction experiment performed on Day 6, mice run in the track for 15 minutes without light or water reward regardless of which port they poked. All behavioral tests were conducted under red lights in the dark cycle of the animal. Behavioral data were analyzed blindly. Please see supplementary information for detailed section on materials and methods.

## Results

### A new behavioral paradigm based on the frustrative non-reward framework

To study FNR, we designed a reward-seeking task that a juvenile mouse can learn within three days. We named this behavioral paradigm “alternate poking reward omission” (APRO). APRO, conducted in a linear track, had two stages which were completed in 5 days (Figure 1A, B). The rate of correct port poking increased over the training period and reached 100% on day 3 (Figure 1C, Table S1). For mice exposed to FNR (FNR mice), frustration was then induced by withholding reward 50% of the time on Day 4 and 80% of the time on Day 5.

**Figure 1.**
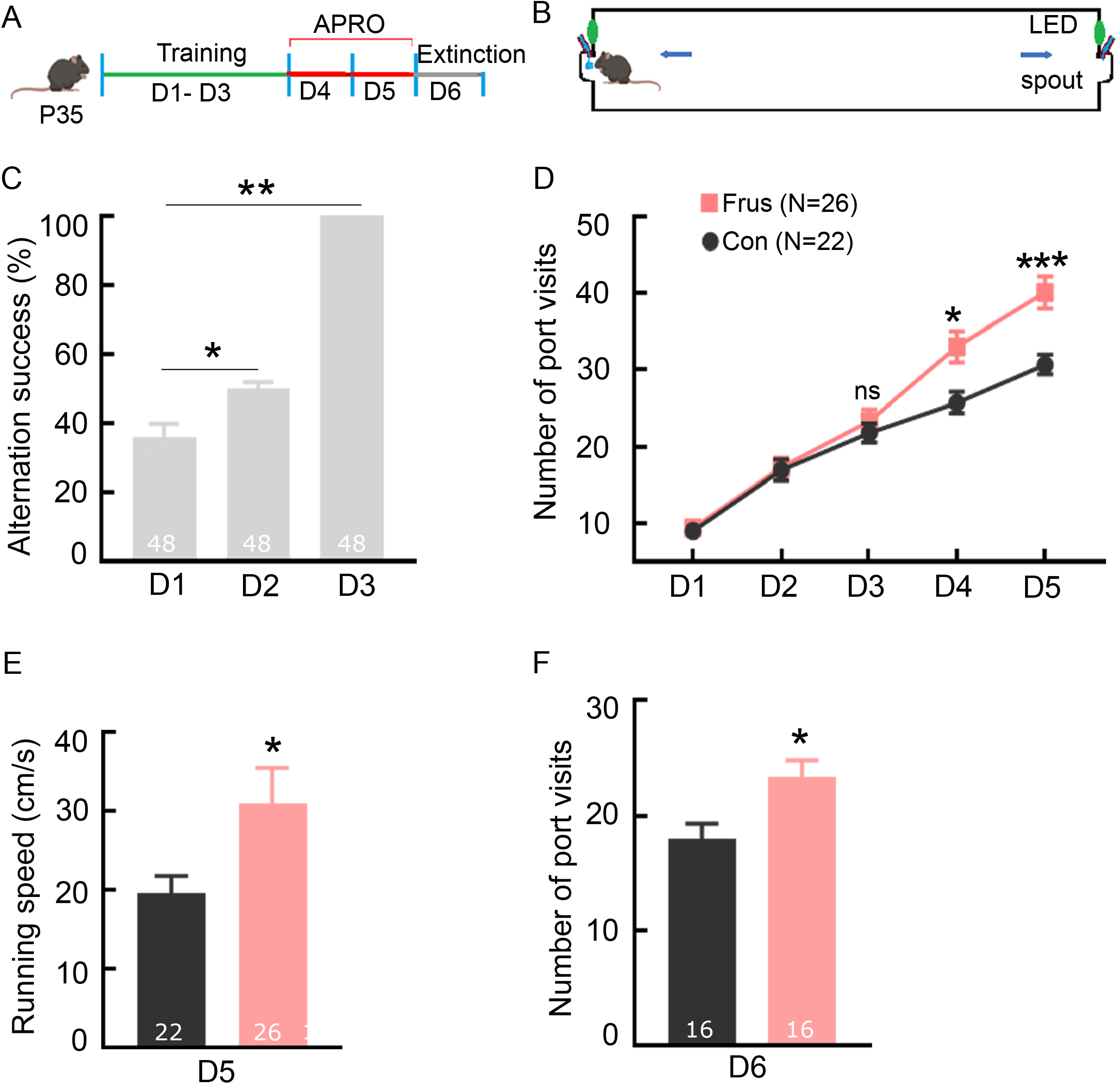
A novel frustrative non-reward paradigm applicable to juvenile mice. (A) Schematic of the experimental paradigm showing the two stages of APRO that begins at P35: training (Days 1-3) and reward omission (FNR; Day 4 & 5). An extinction protocol was applied on Day 6 to test the FNR effect. (B) Schematic drawing of the closed track apparatus installed with infrared-based sensors for poke-triggered cue light and reward delivery system. (C) The percentage of successful alternations between two ports across the three trainings days. (D) The number of port visits in the closed track across days. (E) The average instantaneous running speed between ports on Day 5. (F) The number of port visits during extinction on Day 6. Data are presented as mean + SEM; the numbers in the bars represent the number of animals; * p < 0.05, ** p < 0.01, *** p < 0.001.

FNR mice increased the number of visits to the ports on Days 4 and 5 (Figure 1D, Table S1). Additionally, FNR mice ran faster during the test session than control mice on Day 5 (Figure 1E, Table S1). When testing extinction on Day 6, FNR mice made more visits to the ports than control mice (Figure 1F, Table S1), indicating increased resistance to extinction.

Taken together, the FNR mice’s behaviors in the APRO paradigm are consistent with Amsel’s proposition that frustration increases running speed and resistance to the extinction of instrumental behavior.

### FNR increases locomotion

We assessed the effect of FNR by conducting additional behavioral tests. Frustration is conceptualized as an adaptive motivational state that elevates activity and aggression (22–27). We, therefore, examined locomotion with the open field test (Figure 2A). Mice were tested within 30 min after the test session of APRO on Day 5. FNR mice traveled a longer distance in the arena during the open field test than control mice (Figure 2B, C; Table S1). However, the time spent in the center zone of the arena, which is an indicator of anxiety-like behavior, was comparable in control and FNR mice (Figure 2B, D; Table S1). Both male and female FNR mice exhibited hyperlocomotion (Figure 2B, C; Table S1).

**Figure 2.**
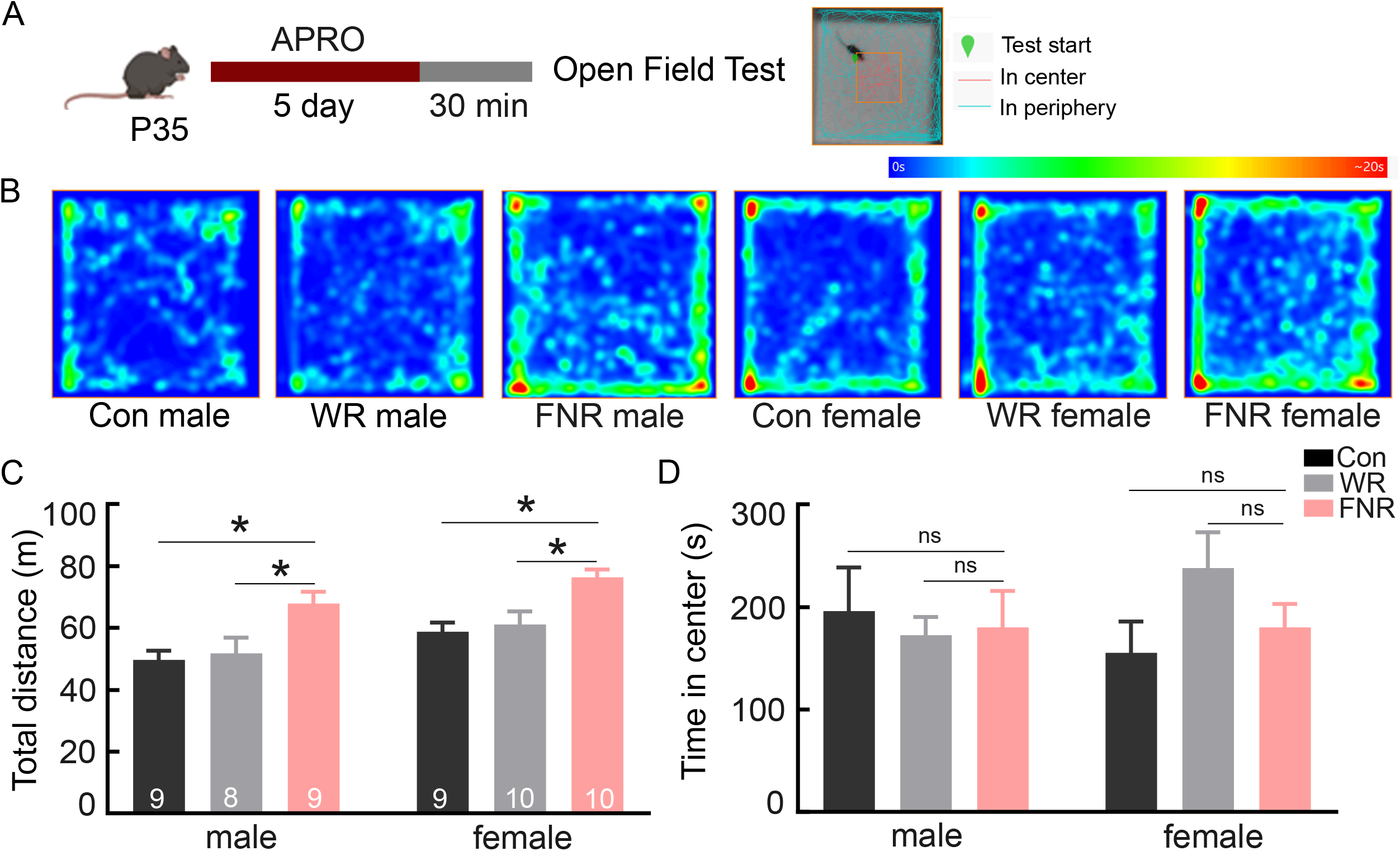
FNR induces hyperlocomotion in both male and female mice. (A) Experimental schedule of APRO followed by open field test and Illustration of division of the central and peripheral zones; the cyan and red lines indicate the mouse’s trajectory in the testing arena. (B) Representative mobility heatmaps showing the activity of male and female mice from different groups within the open field arena. (C) Total distance travelled by control, water restricted, and frustrated mice grouped by sex in a 20-minute testing session. (D) Total time spent in the central zone (25% of the testing arena) by male and female mice of different groups. FNR: frustrative non-reward; WR: water-restricted only; Con: control. Data are presented as mean + SEM; the numbers in the bars represent the number of animals; * p < 0.05.

The mice were under water restriction throughout APRO. Since control mice drank more water than FNR mice in the track prior to the open field test, thirst may underlie the hyperlocomotion of FNR animals. To test the effect of thirst on locomotion, we conducted the open field test in mice who had been water restricted but not exposed to FNR. Water-restricted mice and control mice were comparable in the total distance travelled and time spent in the center zone (Figure 2B, C, D; Table S1). These results indicate that FNR, but not thirst, increases locomotion.

### FNR increases aggression but not non-aggressive social interaction

Next, we examined the effect of FNR on aggression using a modified resident-intruder (RI) test (Figure 3A). A FNR or control mouse (resident) was placed back in the home cage after the test session on Day 5 and allowed to freely explore the cage for 30 min. An unfamiliar, smaller mouse of the same sex (intruder) was then introduced into the same cage to interact with the resident mouse for 5 minutes (Figure 3A).

**Figure 3.**
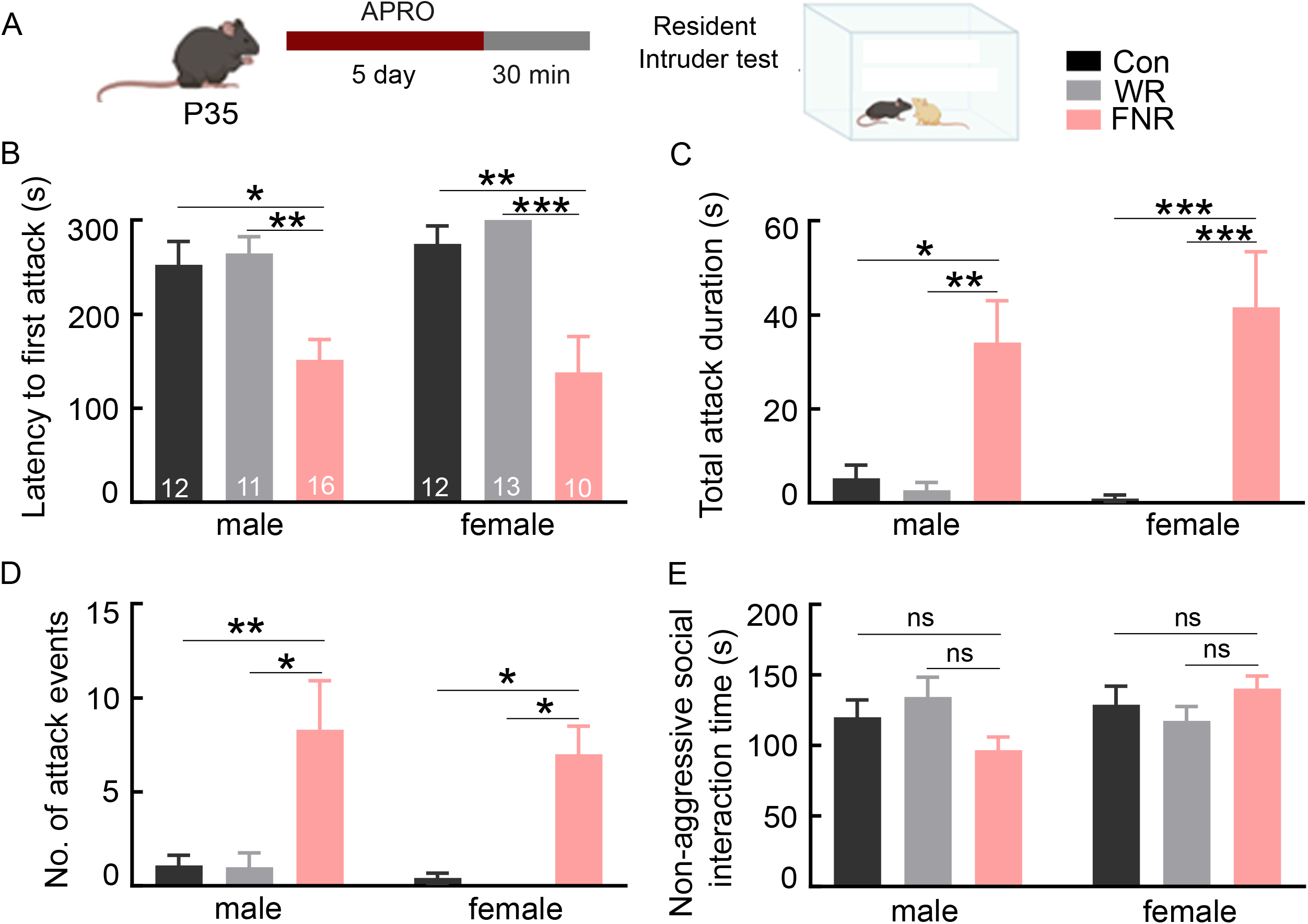
FNR increases aggressive behavior in both male and female mice. (A) Experimental schedule of APRO followed by the resident intruder test. (B) The latency to the first attack in male and female mice. (C) The total time the resident mouse spent attacking the intruder. (D) The number of attack events during the resident-intruder test. (E) The total time spent in non-aggressive social interaction. Data are presented as mean + SEM; the numbers in the bars represent the number of animals; * p < 0.05, ** p < 0.01, *** p < 0.001.

The control mice largely displayed non-aggressive social behaviors such as anogenital sniffing, grooming, and rearing towards the intruder. However, the FNR mice exhibited more aggressive behaviors towards the intruder, such as clinch attack, keep down, lateral threat, and chasing (28–30). Compared to control mice, FNR mice had more total number of attacks, higher average attack duration per episode and total attack time, and a shorter latency to the first attack (Figure 3B-D, Table S1). Notably, C57BL/6 female mice, who usually are not aggressive in the RI test, also attacked the intruder after FNR (Figure 3B-D, Table S1). Water restriction alone had no effect on aggressive behavior (Figure 3B-D, Table S1). Non-aggressive social behavior during the RI test were comparable in FNR and control mice (Figure 3E, Table S1).

We further assessed the effect of FNR on social behavior using the three-chamber sociability test (31) (Figure 4A). FNR and control mice spent similar amounts of time exploring the conspecific and object (Figure 4B, C; Table S1), indicating comparable sociability. Males and females were pooled in data analysis as no sex differences were detected (Table S2). Taken together, these findings indicate that FNR elevates aggression in both male and female mice, without altering non-aggressive social interaction or sociability.

**Figure 4.**
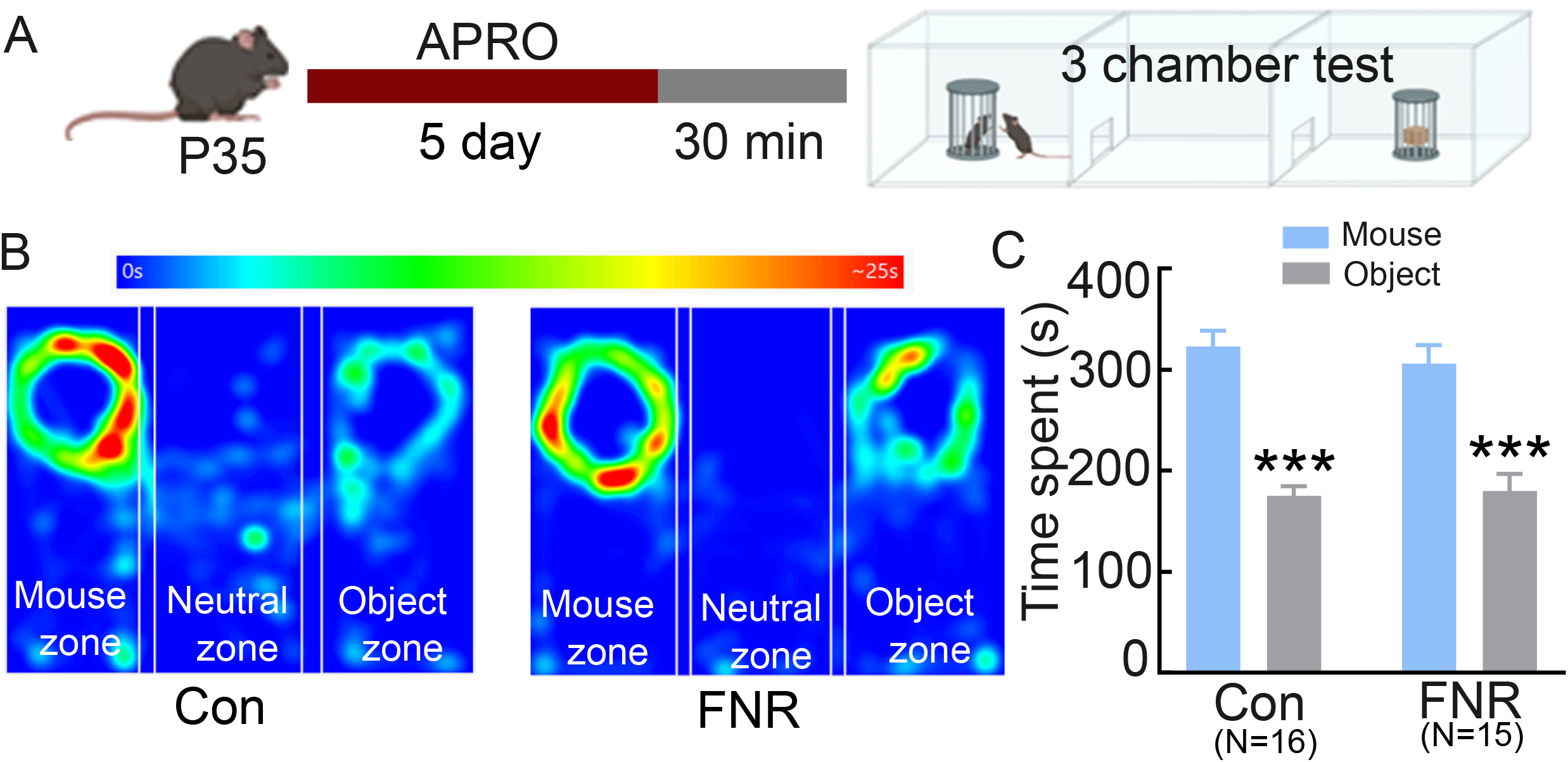
FNR has no effect on social preference. (A) Experimental schedule of APRO followed by the three-chamber social preference test. (B) Representative heat maps showing the activity of the mouse in the mouse and object zone along with the neutral zone. (C)The total time spent with social stimulus (another mouse) and non-social stimulus (object) during the test by control and FNR groups. Data are presented as mean + SEM; N =16 for each group; *** p < 0.001.

### FNR has no effect on anxiety-like or depression-like behaviors

To test whether FNR impacts on anxiety-like behavior, we conducted the light/dark box and elevated zero-maze tests after the test session on Day 5 (Figure 5A). Each test used a different cohort of animals. In the elevated zero-maze test, the time spent in the open arms and the total number of entries into the open arms were comparable in control and FNR mice (Figure 5B, C; Table S1). Similarly, in the light/dark box test, the time spent in the light box and the total number of entries into the light box were comparable between control and FNR mice (Figure 5D, E; Table S1). Males and females were pooled in data analysis as no sex differences were detected (Table S2). These results indicate that FNR has no effect on anxiety-like behavior.

**Figure 5.**
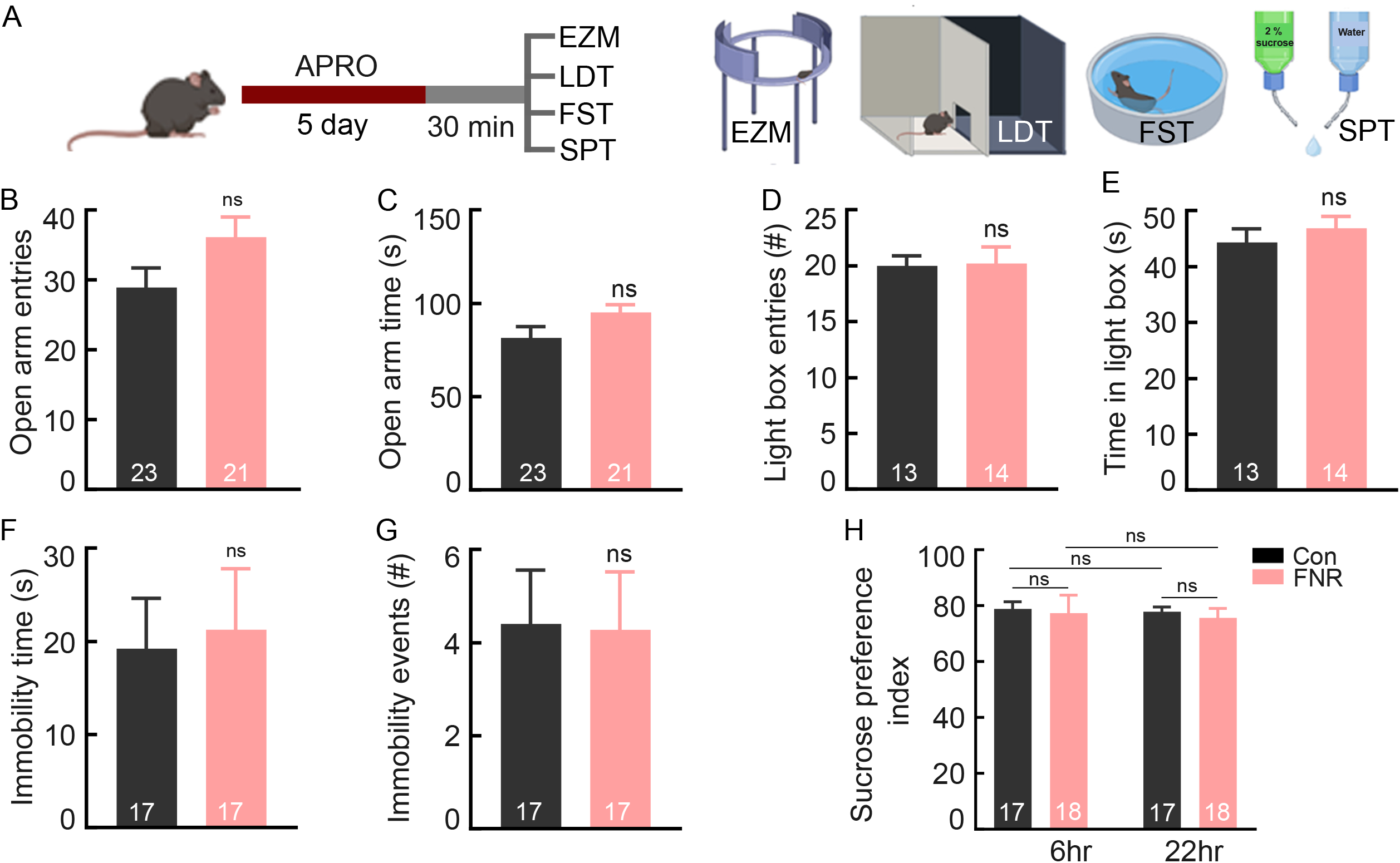
FNR has no effect on anxiety-like or depression-like behaviors. (A) Schematic drawing of the experimental schedule. The elevated zero-maze (EZM), light/dark box test, forced swim test (FST), and sucrose preference test (SPT) followed APRO; different cohorts of mice were used for each test. (B) Total entries of the open arms in the EZM test. (C) Total time spent in the open arms during the EZM test. (D) Total entries of the light box in the light/dark box test. (F) Total time spent in the light box during the light/dark box test. (G) Total time mouse was immobile during FST. (H) The number of immobile events during FST. (H) shows the ratio of sucrose consumption over water recorded 6 and 22 hours after the Day 5 test session in control and FNR group mice. Data are presented as mean + SEM; the numbers in the bars represent the number of animals.

For depression-like behavior, we used the forced swim and sucrose preference tests to assess behavioral despair and anhedonia-like behavior. In the forced swim test, total immobility time and total number of immobile episodes were comparable in control and FNR mice (Figure 5F, G; Table S1). In the sucrose preference test, the ratio of sucrose to water consumed during the 6 hr and 22 hr periods either before or after APRO was comparable in control and FNR mice (Figure 5H, Table S1, S2). Males and females were pooled in data analysis as no sex differences were detected (Table S2).

These results indicate that FNR has no effect on anxiety-like or depression-like behaviors.

### Brain-wide mapping of the effect of FNR on neural activity

To investigate the effect of FNR on brain activity, we removed the brains from the mice (4 control: 2 males and 2 females; 5 FNR: 3 males and 2 females) 90 minutes after the test session on Day 5 to label activated cells expressing c-Fos (Figure S1). The total number of c-Fos positive cells was greater in the FNR than control brains (1.24 ± 0.13×10^6^ for control and 1.95 ± 0.13×10^6^ for FNR; p = 0.001, two-tailed Student’s t-test). We parcellated the whole brain according to the Allen Mouse Brain Common Coordinate Framework and removed regions with only a few c-Fos positive cells from the list for further analysis as the signal-to-noise ratio in these regions was relatively low and the variability between animals was relatively large.

Of the 87 remaining brain regions, 13 regions including prelimbic area, dorsal and ventral parts of the anterior cingulate area, pretectal region, CA2, anterior group of dorsal thalamus, geniculate and medial groups of the dorsal thalamus, cuneiform nucleus, intralaminar nuclei of dorsal thalamus, frontal pole, pons (behavioral state related), and medial region of pallidum showed statistically significant differences between FNR and control mice (two-tailed Student’s t-test corrected for FDR with Benjamini–Hochberg procedure, p < 0.05; Figure 6; Table S3).

**Figure 6.**
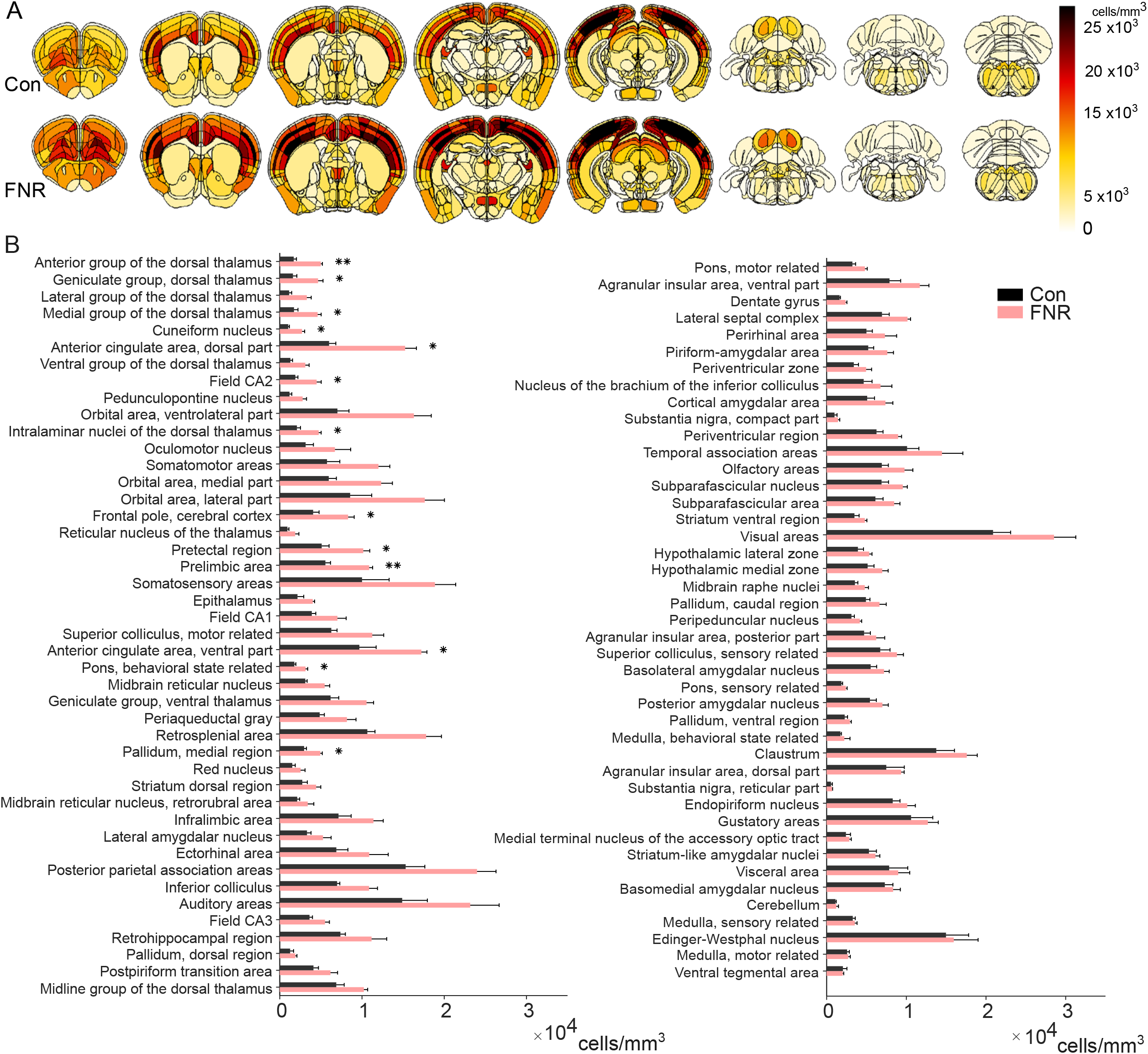
APRO induces brain activation. Control and FNR brains were stained for c-Fos after APRO on Day 5. (A) Heatmaps of the average c-FOS^+^ cell density for the control (n = 4 brains) and FNR (n = 5 brains) groups at different bregma levels. Darker colors indicate higher density of c-FOS^+^ cells. (B) Histograms showing the quantification of c-FOS^+^ cells (cells/mm^3^) in the 87 brain regions; data are presented as mean + SEM; *p<0.05; **p<0.01.

To interrogate the brain response to FNR at a brain-wide level, we constructed correlation matrices for the control and FNR groups (Figure 7A, B). The correlation matrices showed that brain regions within the same anatomical divisions were not always co-activated. Then, based on the correlation matrices, we examined how well the brain functions as an integrated network by computing modularity index Q which quantifies the extent to which the brain can be subdivided into nonoverlapping modules. Modules have a high number of intramodule connections and a low number of intermodule connections (32). High Q indicates that brain modules are organized into a relatively segregated state with many intramodular connections and few intermodular connections, so the brain uses a more localized, segregated processing mode. Low Q indicates that brain modules are organized into a more integrated state with many intermodular connections, so the brain uses a more global processing mode. The FNR brain had lower Q at all network intensities (Figure 7C), suggestive of a more global processing mode than the control brain.

**Figure 7.**
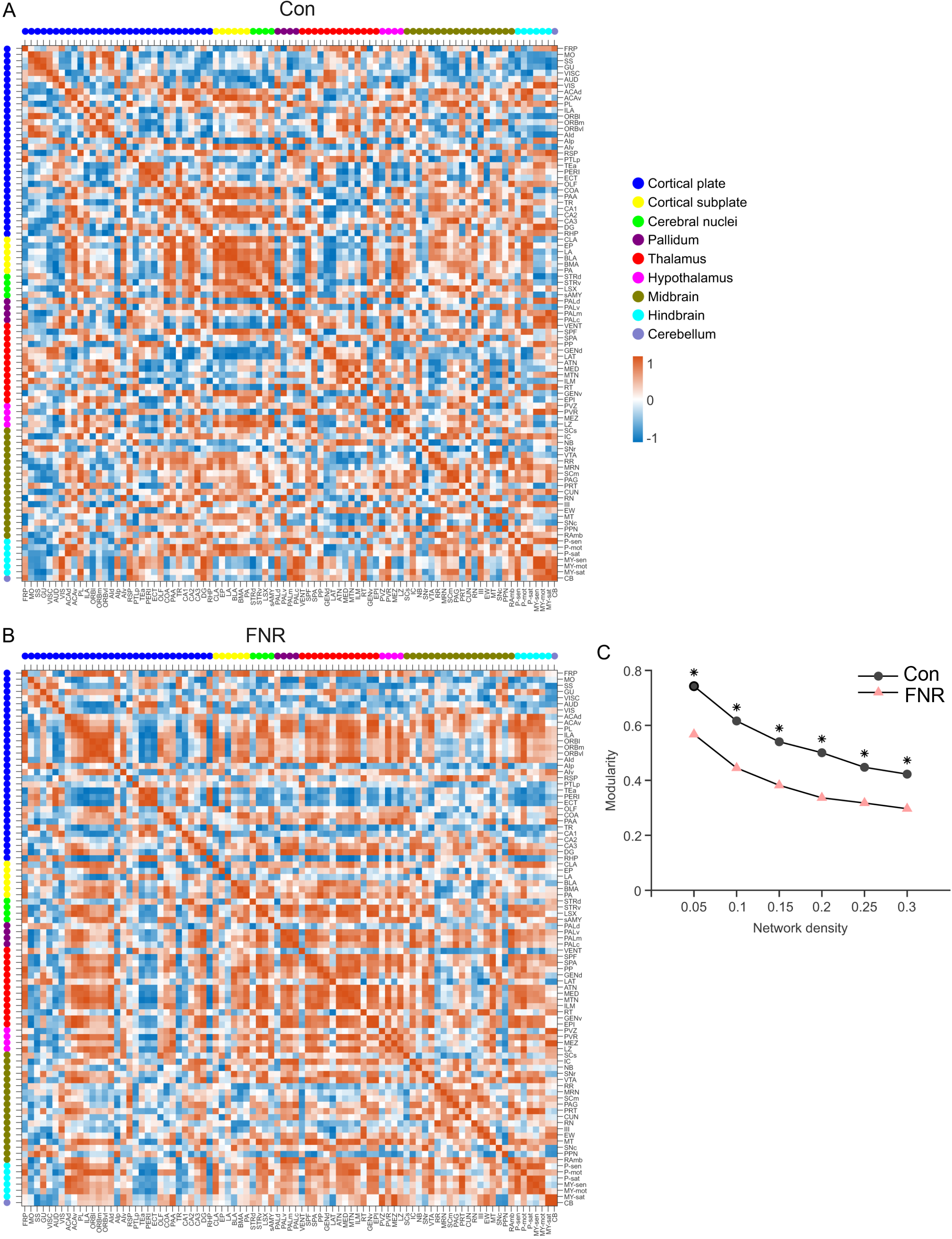
Brain networks of co-activation in control and FNR animals. (A, B) Inter-brain region correlation matrices for control (A) and FNR mice (B). Regions are sorted by anatomical division according to the Allen Brain Atlas. Colored dots at the top and left of the graph represent the anatomical division that each region belongs to. (C) Modularity index across network density thresholding. * p < 0.05.

To compare modular structures between the FNR and control brains, we performed hierarchical consensus clustering analysis (HCC) to build a hierarchy of clusters with increasingly strong interregional correlations (33). HCC revealed a nested community structure with up to 5 levels and 7 modules in the control group, and 6 levels and 7 modules in FNR group. While some brain regions in the same HCC cluster belonged to the same anatomical division, HCC clusters were distinct from the anatomical grouping of brain regions (Figure 7, 8). The cluster composition was different between control and FNR brains (Figure 9), suggestive of a reconfiguration of brain network by FNR. For example, 8 of the 13 brain regions with more c-Fos positive cells in FNR brains were distributed across multiple modules in control brains but clustered in the same module in FNR brains (Figure 9).

**Figure 8.**
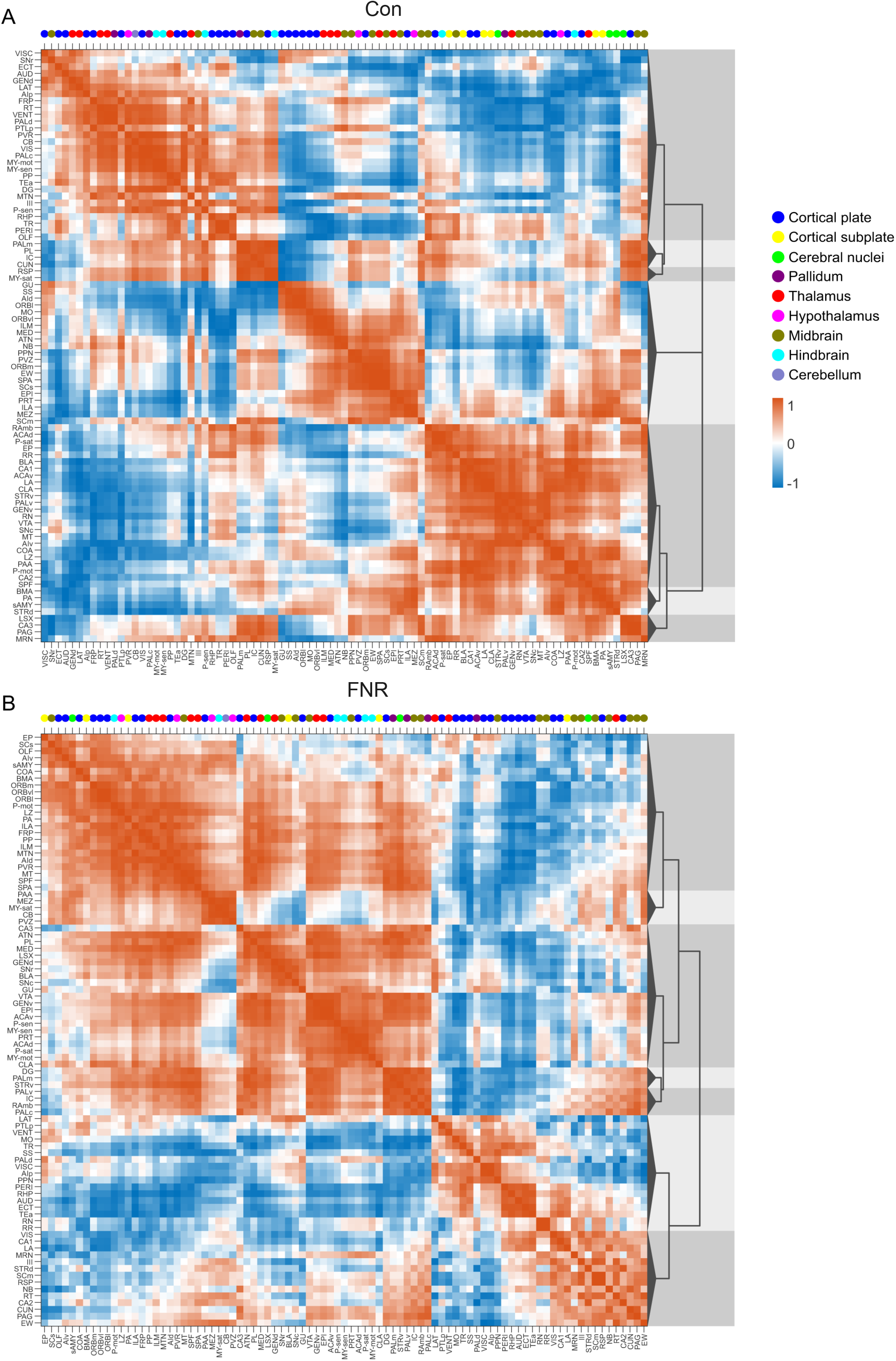
Hierarchical consensus clustering of brain regions by co-activation. Hierarchical consensus clustering of the brain regions for control (A) and FNR mice (B). Regions enclosed by green boxes are highly correlated regions (α < 0.001) The dendrogram to the right of the matrices show the nested modular structures. Colored dots at the top of the matrices represent the anatomical division that each region belongs to.

**Figure 9.**
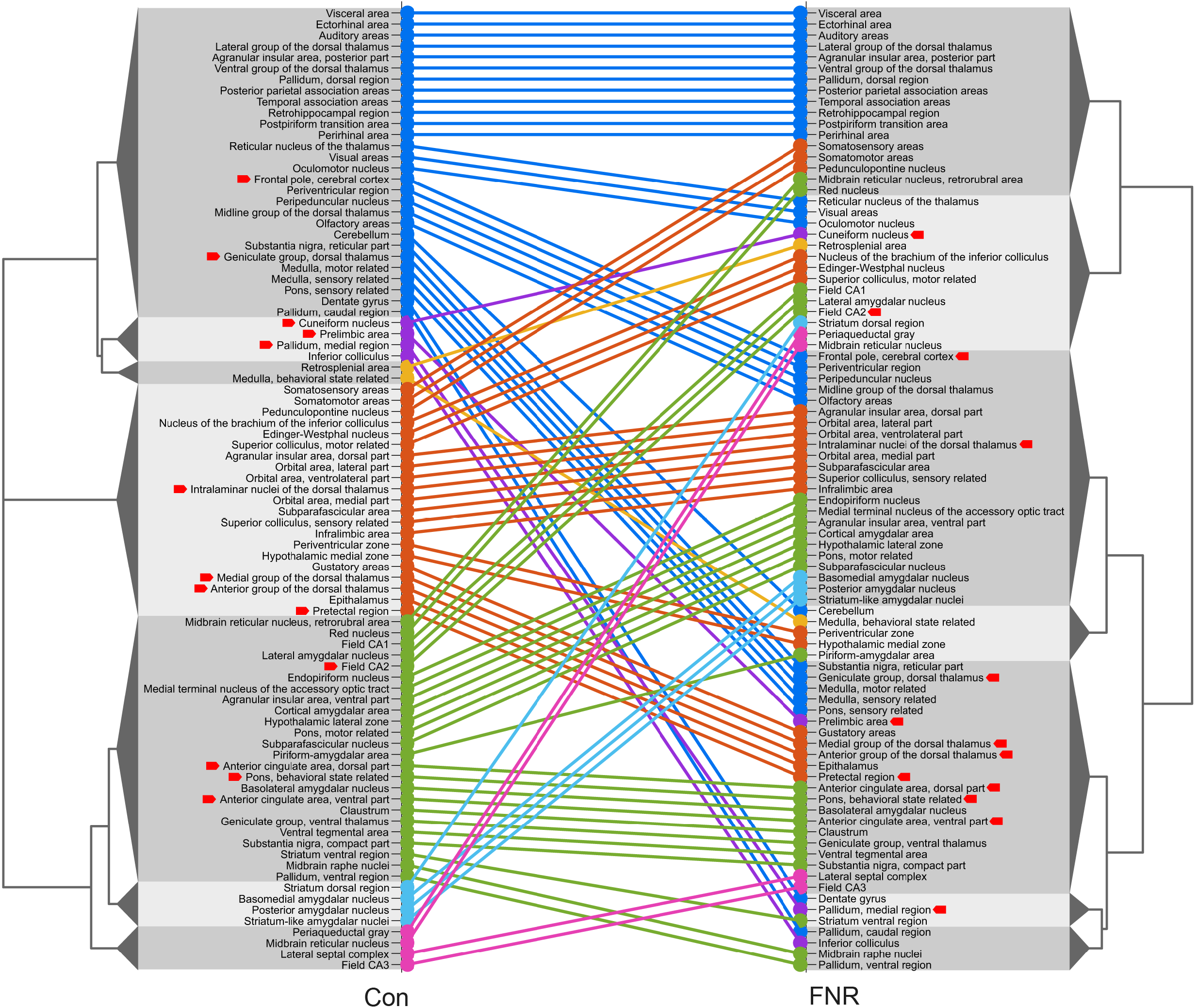
The modular structures of control and FNR brains are different. The same brain regions are connected by lines between the hierarchical consensus clustering dendrograms for control (left) and FNR (right) brains. Brain regions within the same cluster of the control brain are indicated with the same color and the same color scheme applied to both control and FNR brains. Red arrows point to the brain regions that show significant differences in c-Fos density between control and FNR mice.

These results indicate that FNR increases neural activity in selective brain regions and changes the processing mode of brain networks.

## Discussion

Irritability is among the most common reasons that children are brought for psychiatric evaluation and care (1, 2, 6, 34, 35). Despite the significant impact of irritability on mental health, little is known about its underlying neural mechanisms and limited treatment options are available (4, 36). Mechanistic understanding of irritability is needed to guide development of new treatments. Youths with severe irritability have elevated responses to FNR; the latter is a cross-species, translational construct in the negative valance system of Research Domain Criteria (RDoC) (9, 36, 37). Therefore, translational research elucidating the neurobiology of FNR can facilitate research on the pathophysiology of irritability. Importantly, even though Amsel’s frustration theory has been examined extensively from a behavioral perspective since the 1950s, few studies attempt to disentangle the underlying brain circuits and neural mechanisms.

To that end, we developed a mouse behavioral paradigm (APRO) based on FNR to study irritability-like behavior in juvenile mice. This paradigm takes advantage of the mouse’s natural tendency of alternating visits between two places to complete training within 3 days, significantly shorter than older paradigms e.g., double runway and operant conditioning (14–16). We demonstrated that following APRO, FNR mice increased locomotion, aggression, and resistance to extinction of learned behavior. These behavioral alterations are consistent with frustration effects reported in rats, chimpanzees, pigeons, and humans (10, 11, 13, 22–24, 26, 38–42). However, our FNR mice had no change in anxiety-like or depression-like behavior, or social preference. These findings suggests that APRO induces relatively specific behavioral changes that are directly relevant to those seen in youth with severe irritability who have frequent temper outbursts, characterized by increased motor activity and aggression (42, 43). It is noted that APRO has more selective behavioral effects than frustration induced by extinction tasks that completely omit reward (44).

Functional magnetic resonance imaging (fMRI) based on blood-oxygenation−dependent signals (BOLD) has been used to study brain activity related to irritability in humans. However, the spatial resolution, signal to noise ratio, and incomplete understanding of the factors contributing to BOLD signals limit the precision and details of the derived brain activity map. We elected to assess brain activity from c-Fos because c-Fos is a commonly used marker of cell activation (45–48) and its nuclear localization assists machine learning-based cell segmentation and automated cell counting. Whole-brain imaging of c-Fos labeling provides more refined and direct measurement of neural activity compared to fMRI studies.

13 brain regions including prelimbic area, dorsal and ventral parts of the anterior cingulate area, pretectal region, CA2, anterior group of dorsal thalamus, geniculate and medial groups of the dorsal thalamus, cuneiform nucleus, intralaminar nuclei of dorsal thalamus, frontal pole, pons (behavioral state related), and medial region of pallidum showed significant c-Fos increase in FNR, compared to control, mice. Some of them are within the limbic system and have been associated with negative emotions and high irritability, such as the hippocampus, thalamus, and ACC. The hippocampus plays a role in learning, memory, and emotional processing. ACC lesions in humans are associated with hostility, emotional unconcern, and irresponsibility (24, 49). Besides being a sensory relay center, the role of the anterior and midline group thalamic nuclei in response to aversive and rewarding stimuli has emerged from recent rodent studies (50–53). In addition to individual brain regions, we explored the effect of FNR on the brain network and found that FNR changes its modular structures, specifically reducing modularity and altering the modular composition, so that brain regions recluster and more regions work together in a cooperative and integrated manner. This is consistent with the fMRI finding that FNR changes the degree of brain network segregation and reconfigures brain modules in humans (9). The brain regions and modules altered by FNR can be potentially involved in the behavioral outcomes of FNR. Their exact functions await to be delineated by future investigations with region-specific manipulation of neural activities. Our study generated a candidate brain region list that serves as a starting point and raises many intriguing questions for such investigations.

Our APRO paradigm produces robust and specific behavioral effects that recapitulate the behavioral phenotypes of DMDD. The full capacity of this novel paradigm in studying irritability has yet to be determined by more comprehensive testing in future, such as if the frustration effect is strain-specific, how long the frustration effect lasts, and if there are other behavioral domains that are also affected. It is possible that APRO produces strain specific behavioral changes given the genetic, physiological, and behavioral variations among mouse strains. If this is true, research on strain-dependent responses to APRO would facilitate clinical research on individual differences in irritability. Applying APRO to mice predisposed to anxiety-like or depression-like behavior, such as mice harboring mutations in genes increasing risk for psychiatric disorders or exposed to environmental risk factors, is another avenue of future research that could provide insights into the complex pathogenic mechanisms of human irritability.

c-Fos labeling is a robust method to generate a whole-brain activation map, which guides the formulation of hypotheses for further studies. As the animal is typically perfused ~90 min post stimulation to allow c-Fos proteins to accumulate to detectable levels (45), the temporal resolution of c-Fos reported brain activation is low. The fixation step also makes this method unsuitable for time-lapse imaging in the same subject. In addition, the specific type of neurons that are activated are unclear from measurement of endogenous c-Fos as it is ubiquitously expressed in all neurons. This limitation can be addressed in future studies using transgenic mice that express c-Fos promoter driven fluorescent proteins in a cell-type specific manner. We did not detect sex differences in the brain-wide c-Fos imaging study. This is consistent with the behavioral results. Although the relevant studies are relatively small and non-representative, currently there is no strong clinical evidence supporting gender differences in the pathophysiology of irritability. However, our c-Fos study could be underpowered to detect sex differences due to the small sample size.

Despite the aforementioned limitations, our study provides a novel behavioral paradigm for studying FNR in juvenile mice and a landscape of brain activity changes in response to FNR. This study lays a foundation for further research on the neural mechanism of FNR and irritability by precisely manipulating and measuring neural activity with advanced genetic, optogenetic, and electrophysiological tools.

## Acknowledgements and Funding

This work was supported by the Intramural Research Program of the National Institute of Mental Health (ZIAMH002881 to Z.L. and ZIAMH002786 to E.L.) and the Center on Compulsive Behaviors (CCB), NIH via the NIH Shared resource Subcommittee to A.N. We thank Dr. Ted Usdin (NIMH) for co-mentoring A.N. for the CCB fellowship program and summer student Jeffrey Zhang from Oberlin College and Conservatory for assistance in behavioral data analysis. We thank NIMH IRP Rodent Behavior core for support of behavioral analysis and LifeCanvas Technologies for assistance with c-Fos whole brain immunohistochemistry and light-sheet imaging.

## Supplementary Information

### METHODS AND MATERIALS

#### Animals

All animal procedures followed the US National Institutes of Health Guidelines Using Animals in Intramural Research and were approved by the National Institute of Mental Health Animal Care and Use Committee. C57BL/6J mice and BALB/c mice used in this study were bred in house using breeders purchased from The Jackson Laboratory (#000664, #000651). Mice used in behavioral experiments were maintained under a 12-hour reversed light (8 pm–8 am)/dark (8 am–8 pm) cycle with access to water and food *ad libitum* except during the APRO period when animals were given ad libitum access to water for 1 hr per day. Mice were individually housed for two days before aggression testing. For all other behavioral tests, two mice were housed in a divided cage and separated by a partition. Animals were randomly assigned to experimental groups.

#### APRO

Mice (35 days old; male and female) were placed under water restriction for three days during which they were given ad libitum access to water for 1 hour per day. The apparatus used for APRO was constructed by NIMH Section on Instrumentation. It is a closed running track (50L x 10W x 15H cm) with a spout installed at each end and a LED light mounted next to the spout. The mouse was allowed to run freely in the track for one 15-min session per day. The mouse could lick the spout when it reached the end of the track. If the mouse licked the spout other than the previous one it had licked, the LED light came on for 2 sec and a water drop was delivered from the spout.

During training sessions, the mouse received water reward for every correct spout licking. Only mice making ≥12 licks on each side by Day 3 proceeded to Day 4 and 5 of APRO. On Days 4 and 5, mice run on the same track with the same rules. While control mice received water reward for all correct lickings on Days 4 and 5, water reward was delivered for only 50% of correct lickings on Day 4 and 20% on Day 5 for mice subjected to FNR. Thirty minutes after the FNR session on Day 5, the mice were subjected to a battery of behavioral tests, including the open field, resident intruder, elevated zero-maze, light/dark, forced swim, sucrose preference, and three-chamber social preference tests. Different cohorts of mice were used for each behavioral test so that each mouse was subjected to only one behavioral test. For the extinction experiment performed on Day 6, mice were run in the track for 15 minutes without light or water reward regardless of which spout they licked. All behavioral tests were conducted under red lights in the dark cycle of the animal. Behavioral data were analyzed blindly.

#### Behavioral testing

##### Open field test

Mice were allowed to freely explore an opaque box (48L x 48W x 40H cm) for 20 minutes. The behavioral data were analyzed by using ANY-maze software (Stoelting, USA) to calculate the total distance travelled, and the time spent in the center zone that accounted for 25% of the total area.

##### Resident Intruder Test

After the training session on Day 3, mice were housed singly in cages. Resident Intruder tests were conducted in the animal’s home cage. On Day 5, mice were returned to their home cages after the FNR session. A smaller BALB/c mouse of the same sex was introduced into the cage 30 min later. The cage was covered with a transparent plexiglass lid. The two mice were allowed to interact for 5 min. If bodily injury resulted from the interaction between the resident and intruder mice, testing was discontinued, and the data were excluded from further analysis. The data from the resident intruder test were analyzed manually by a researcher blinded to experimental conditions. Based on previous studies, clinching, chasing, boxing, pinning, keeping down, and wrestling were identified as aggressive behavior (1, 2). Anogenital sniffing, grooming, inquiry, and flank rubbing were classified as nonaggressive social behaviors (3).

##### Elevated zero-maze test

Mice were allowed to freely explore an elevated zero-maze (blue acrylic annular platform, 105 cm in diameter and 5.5 cm wide, elevated 60 cm above the ground) for 5 minutes. The maze consisted of two closed quadrants with walls of 30-cm high and two open quadrants with no walls. ANY-maze software was used to calculate time spent, and entries to, the open quadrant.

##### Light/dark box test

The test box was divided into two compartments (27 L x 27 W x 30 H cm) with an opening in between to allow the mouse to enter. The light chamber had white walls and was illuminated, while the dark compartment had black walls and was not illuminated. Mice were placed in the light box when the test started and allowed to freely explore the box for 5 min. ANY-maze software was used to calculate the percentage of time spent in the light compartment and the number of transitions between the light and dark compartments.

##### Forced swim test

Mice were gently released into a Plexiglas cylinder (20 H x 10 D cm) filled with water (23°C, 7.5 cm deep) and left in the water for 5 minutes. ANY-maze software was used to calculate the total immobility time, number of immobile episodes, and delay to the first immobile episode.

##### Three-chamber social preference test

The social preference test described by Rein et al., 2020 (4) was used with some modifications. The test has three phases: habituation, pre-testing, and testing. During habituation, the openings between the central and side chambers were blocked. There was an empty metal wire cup inside each side chamber. The mouse was placed in the central chamber and allowed to freely explore for 5 min. During the pre-test, the blocks between the central and side chambers were removed. Identical objects (blue Lego bricks) were placed in the cups. The mouse was allowed to interact freely with the objects for 5 min. After the pre-testing phase, the mouse was held temporarily in the central chamber. One object was replaced by a social stimulus (an age- and sex-matched C57BL/6 mouse) and another object was replaced by a novel object (hexagonal Lego brick). The mouse was then released to freely explore the social and non-social stimuli for 10 min. ANY-maze software was used to record the time spent with social stimulus (mouse) and non-social stimulus (object) during the test session.

##### Sucrose preference test

The sucrose preference test was performed as described by Liu et al., 2018 (5), with modifications. The test has three phases: adaptation, baseline measurement, and testing after FNR. During the adaptation phase, two bottles, one containing water and the other one containing 2% sucrose solution, were inserted into the cage lid. The mouse (P29) was housed in a cage with two bottles for 2 days. The position of the bottles was switched on the second day. Both bottles were removed at 6 pm of the second day. The baseline measurement started at 9 am of the following day. Fresh bottles containing 2% sucrose and water were weighed and inserted into the cage lid. The bottles were weighed 6 hrs and 22 hrs later to determine the baseline sucrose preference index. Mice who consumed no sucrose solution during the baseline measurement period were excluded from further testing. The sucrose preference testing after FNR started 1 hr after the animal was returned to its home cage following the FNR session. Weighed water bottle and sucrose solution bottle were inserted into the cage lid. The bottles were weighed 6 hr and 22 hr later to determine the post-FNR sucrose preference index.

#### c-Fos imaging

##### Brain clearing, immunohistochemistry, light sheet microscopy, and image analysis

Mice anesthetized with isoflurane were transcardially perfused with cold phosphate buffered saline (PBS, PH 7.4) and 4% paraformaldehyde (PFA) in PBS sequentially. The brain was removed and post-fixed in PFA for 24 hrs at 4°C, followed by rinsing in PBS three times (30 min each time) before being transferred to PBS containing 0.1% sodium azide. Brain clearing, staining, imaging, and image analysis were conducted by LifeCanvas Technologies. The brain was cleared using the SHEILD method and immunolabelled using eFLASH technology with stochastic electrotransport and SWITCH. The brains were cleared in the delipidation buffer at 42°C for 48 hr using Smartpro Clear II. Cleared brains were co-incubated with the rabbit anti-c-Fos monoclonal antibody (Abcam, #ab214672) and mouse anti-NeuN monoclonal antibody (Encor Biotechnology, #MCA-1B7). After rinse, the brains were incubated with secondary antibodies in ratios of 1:2 (primary:secondary) molar ratio (Jackson ImmunoResearch).

After immunostaining, the brains were incubated with EasyIndex to render them transparent (RI = 1.5). The brains were then mounted in agar blocks, re-immersed in EasyIndex to match the refractive index of the brain and imaged with a SmartSPIM light sheet microscope. Images were acquired throughout the brain at 3.6X magnification, 1.8 µm/pixel resolution on the XY plane, and 4 µm thickness along the z-axis. After image acquisition, the images were tile-corrected, de-striped, and registered to the Allen Brain Atlas using an automated process (6).

A custom convolutional neural network was used for automated detection of c-Fos positive cells using the TensorFlow python package. Two networks were used: a U-Net architecture based fully-convolutional network was used for detection and a ResNet based convolutional network was used to classify a c-FOS positive or negative hit (7, 8). The c-Fos images were registered with the Allen Brain Atlas to determine the location of c-FOS positive cells. The density of c-Fos positive cells for each brain region were calculated by dividing the total number of c-Fos positive cells in each brain region by the region volume, then averaged between the two hemispheres.

##### Construction of brain co-activation networks

We constructed connected, undirected, signed, weighted networks for the control and FNR groups from interregional correlations for the control and FNR groups by Pearson correlation coefficients. To reduce the high Pearson correlation coefficients caused by system variability from brain to brain, the density of c-Fos positive neurons was normalized by multiplying the density in 87 brain regions in the same brain by a same factor to make the total number of c-Fos positive neurons in each brain equal the mean within the same treatment group (control or FNR). Pearson correlation coefficient was calculated between all pairs of the 87 brain regions within the same treatment group.

##### Calculation of modularity index

To compare network properties between the control and FNR groups, we calculated the modularity index Q, which quantifies the extent to which the brain exhibits a modular structure. The modularity index was calculated using the Louvain greedy algorithm (9) implemented in the Brain Connectivity Toolbox (10). We set the threshold of network density to 0.05, 0.1, 0.15, 0.2, 0.25 and 0.3. Network connections below threshold were removed. The Louvain algorithm is a heuristic to detect communities based on maximizing the modularity index. First, the algorithm combines nodes into small communities by maximizing modularity locally. Second, the small communities were used as new nodes to build a new network. This process is repeated until there is no further increase in modularity (9–11). Considering the stochastic initialization of the greedy optimization, we applied the algorithm 1000 times for each group, and used the highest modularity index. To calculate modularity, we used vector *g* to denote partition of a network, where *g_i_* denoted the cluster assignment of node *i*. The following modularity index formula adapted from Reichardt and Bornholdt (12) was used:

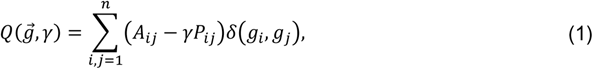

where *A* is the connection matrix, *P* is the expected connection matrix under a null model, *γ* is the resolution parameter, and *δ (g_i_, g_j_)* is the community co-assignment matrix, where *δ* = 1 if *g_i_* = *g_j_*, and *δ* = 0 otherwise. We used the following typical null model to derive *P*:

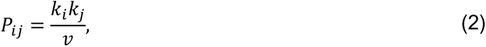

where *k* was the sum of all correlations of a given node, and *v* was the sum of all *k* values. We set *γ* to 1 for both control and FNR groups. To estimate the statistical significance of the difference in modularity index between groups, permutation tests were used by shuffling the relationship between animal ID and c-Fos positive neuron counts within each brain region. 5000 permutations were used. The permutation test results for each network density were then corrected for multiple comparisons by estimating false discovery rate (FDR) with Benjamini–Hochberg procedure.

##### Data and statistical analysis

Graphpad prism (9.5.1) was utilized for statistical analysis. For behavioral analysis, two-tailed Student’s t-test was used for normally distributed data with equal variance and Mann-Whitney U test was used for data that did not satisfy these requirements. One-way ANOVA on ranks or two-way ANOVA with Dunn’s procedures or Tukey’s test for post hoc multiple comparisons were used for comparing more than two groups. The density of c-Fos positive neurons was analyzed with two-tailed Student’s t-test to compare between control and FNR mice and the obtained p-values were adjusted with the FDR approach. P < 0.05 was considered significant. All data are presented as the mean + SEM. Statistical results for all figures are provided in Table S1.

---

**Fig. S1.**
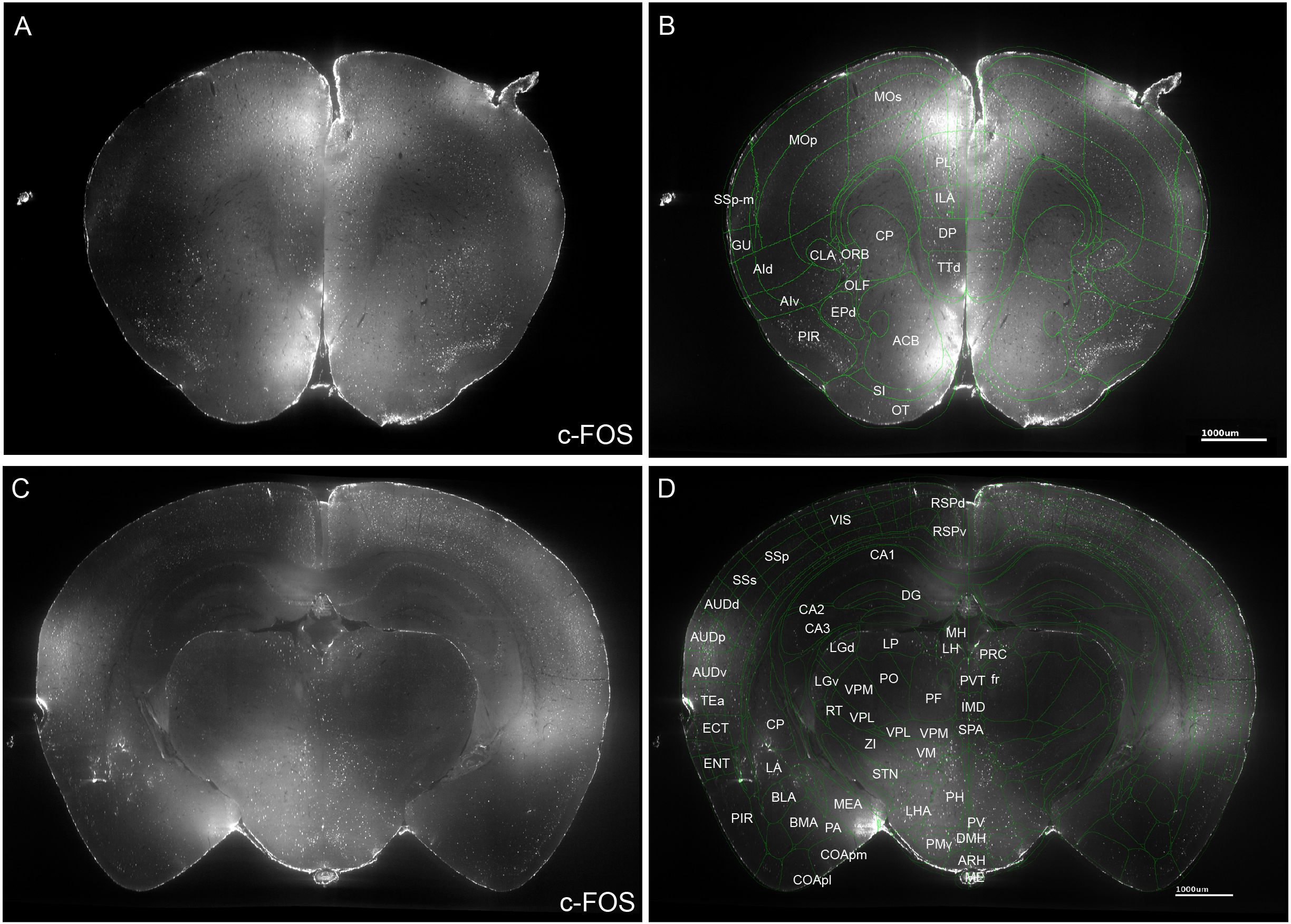
Representative c-Fos images and registration to Allen Brain Atlas. The brains were processed for c-Fos staining after APRO and imaged with light-sheet microscopy. (A) Representative image taken at the level of the medial prefrontal cortex. (B) The image in A was registered with the Allen Brain Atlas and the 87 brain regions localized in this image were labeled. (C) Representative image containing the hippocampus. (D) The image in C was registered with the Allen Brain Atlas and the 87 brain regions localized in this image were labeled.

